# Data-driven workflow for comprehensive gene expression analysis in complex microbiomes

**DOI:** 10.1101/2025.01.17.632662

**Authors:** Ryo Mameda, Hidemasa Bono

**Author notes:** **Correspondence:** Hidemasa Bono.

## Abstract

Complex microbiomes, characterized by highly diverse microbial species, are ubiquitous in the environment. However, the genomes of many constituent microbes remain uncharacterized, complicating the use of publicly available genome sequences as reference data for gene expression analysis based on next-generation sequencing (NGS) data. Consequently, selecting suitable reference sequences is critical for accurate gene expression analysis in complex microbiomes. Additionally, such analyses require precise quantitative evaluation of the genes within each microorganism. In this study, we identified appropriate reference sequences and developed a data-driven analytical workflow for gene expression analysis in complex microbiomes. Utilizing metagenomic and metatranscriptomic NGS reads from these microbiomes, the workflow evaluates gene abundance and transcriptional levels, addressing key analytical challenges. The findings demonstrate that metagenomic contigs are more suitable than conventional reference sequences for mapping both metagenomic and metatranscriptomic data. Moreover, this workflow facilitates not only the assessment of transcriptional activity but also the evaluation of gene expression potential.

## Introduction

Complex microbiomes, such as soil microbes, aquatic microbes, and gut bacteria, are ubiquitous in the environment and play critical roles in influencing the higher organisms with which they coexist. Additionally, certain complex microbiomes, such as those in activated sludge, are utilized in industrial applications, making the elucidation of their functions highly valuable.

Metagenomic and metatranscriptomic analyses using next-generation sequencing (NGS) have been widely employed to investigate the functions of complex microbiomes. Studies targeting individual microbial species include investigations of nitrification genes in Nitrosomonas sp. within activated sludge [1] and the discovery of antibiotic biosynthesis genes in *Actinomycetes* from soil microbiomes [2]. Broader processes, such as the nitrogen cycle [3] and carbon cycle [4], have also been analyzed in soil microbiomes. These examples underscore the increasing application of metagenomic and metatranscriptomic approaches in functional microbiome research.

Selecting appropriate reference sequences is a critical factor in the functional analysis of complex microbiomes. In NGS-based analyses, NGS reads must be compared against reference sequences with functional annotations. For single organisms, complete genome sequences can serve as references; however, this approach is challenging for complex environmental microbiomes due to their high diversity. Recent advancements in microbial genome sequencing have introduced methods such as metagenome-assembled genomes (MAGs) for bacteria with relative abundances of less than 1% in complex microbiomes [5] and single-amplified genomes (SAGs) obtained through single-cell analysis [6]. Notably, approximately half of the genomes in Release 220 of the Genome Taxonomy Database consist of MAGs or SAGs [7]. Despite these advances, only an estimated 2.1% of environmental bacteria have been genome-sequenced [8], highlighting the limitations of using database-registered genome data as references for species and gene function identification in complex microbiomes. Comprehensive functional analyses of complex microbiomes require approaches that encompass the entire microbial community, regardless of genome sequencing status. Thus, selecting effective reference sequences is paramount.

In addition to reference selection, methods for evaluating gene expression levels must also be considered. Distinguishing between microbes lacking target genes and those possessing but not expressing them is critical. For example, a study on nitrifying bacteria in activated sludge reported that genes such as nitrate reductase exhibited extremely low RNA levels relative to their DNA abundance, suggesting the genes possess only the potential for expression [9]. This underscores the importance of analyzing both DNA and RNA from complex microbiomes to enable advanced gene expression studies. Such analyses necessitate reference sequences for both DNA- and RNA-derived NGS reads.

To date, several proposed approaches for gene expression analysis in complex microbiomes primarily rely on metatranscriptomic reads or MAGs [10–12]. However, metatranscriptomic reads are limited to evaluating actively expressed genes, while MAGs often lack a substantial portion of genes present in microbial communities [13]. Consequently, there is an increasing demand for workflows capable of comprehensively analyzing the presence of functional genes in metagenomes, their expression status, and variations in gene expression levels across samples.

This study aimed to develop a functional analysis workflow for complex microbiomes using metagenomic and metatranscriptomic reads from the same samples available in the NCBI database.

## Methods

### Computational resources

This analysis was conducted using a system equipped with 64 GB RAM and an Apple M1 Max chip, operating on macOS Sequoia. The binning step was performed on a Linux environment using the NIG supercomputer at the ROIS National Institute of Genetics.

### Reference sequence analysis

The analysis utilized 56 short-read datasets of soil microbiomes. Metagenomic and metatranscriptomic reads were obtained from 24 samples each, as described in reference studies [14–17] (Supplementary Table 1). FASTQ files for these reads were retrieved from the NCBI Sequence Read Archive database using the SRA Toolkit [18] (v3.0.10) with the prefetch and fasterq-dump commands. Quality control and trimming were performed using fastp [19] (v0.23.4) with the parameters -q 20 -t 1 -T 1. Three types of candidate reference sequences were evaluated to optimize reference selection for complex microbial gene expression analysis. Trimmed metagenomic reads were assembled into contigs using MEGAHIT [20,21] (v1.2.6) with default parameters. The assembled contigs were binned using MetaBAT2 [22] (v2.15) and MaxBin2 [23] (v2.2.7), and the quality of the resulting bins was assessed using CheckM [24–26] (v1.1.11). Binned contigs with ≥50% completeness and ≤25% contamination were designated as MAGs. To construct metatranscriptomic contigs, trimmed metatranscriptomic reads were assembled using Trinity [27] (v2.15.1) with default parameters. Quality checks for metagenomic contigs, MAGs, and metatranscriptomic contigs were performed using SeqKit [28,29] (v2.8.0) with the stats command. Protein-coding sequences were predicted from all candidate reference sequences using Prodigal [26] (v2.6.3) with the -p meta parameter. Trimmed metagenomic and metatranscriptomic reads were mapped to the predicted protein-coding sequences using BWA MEM [30] (v0.7.17) or Bowtie2 [31–33] (v2.5.1) with the --sensitive parameter. SAM files generated during mapping were converted to BAM format using SAMtools [34] (v1.17) with the sort command, and mapping statistics were analyzed with the flagstat command.

### Gene expression analysis

Gene annotation was assigned to the predicted protein-coding sequences as follows. Predicted nucleotide sequences of metagenomic contigs were screened against rRNA sequences from NCBI [35] using BLASTN [36] (v2.15.0) with an E-value threshold of 0.1; sequences matching rRNA were excluded. Remaining protein sequences were annotated by querying against the Swiss-Prot database of well-characterized proteins in UniProt [37] using DIAMOND [38] (v2.0.15) with BLASTP and an E-value threshold of 0.1. Sequences without hits in Swiss-Prot were further annotated using protein domain information from Pfam [39] via HMMER [40] (v3.3.2) with an E-value threshold of 0.1, and were parallelized using GNU Parallel [41]. Predicted protein sequences were annotated based on the highest-ranking hits from the BLASTP and HMMER searches, with sequences lacking significant matches designated as hypothetical proteins. Using the annotations and BAM files from the previous steps, read counts for each sequence were quantified with the featureCounts command in Subread [42] (v2.0.6) (Fig. 1). Both metagenomic and metatranscriptomic read counts were normalized using the following equations: for a given contig *t*, let *G*_*t*_ represent the metagenomic read count, *T*_*t*_ the metatranscriptomic read count, and *L*_*t*_ the contig length. Genes per million (GPMt) and transcripts per million (TPMt) were calculated accordingly.

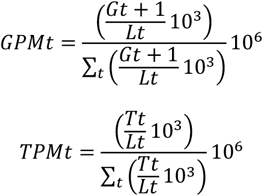

**Figure 1.**
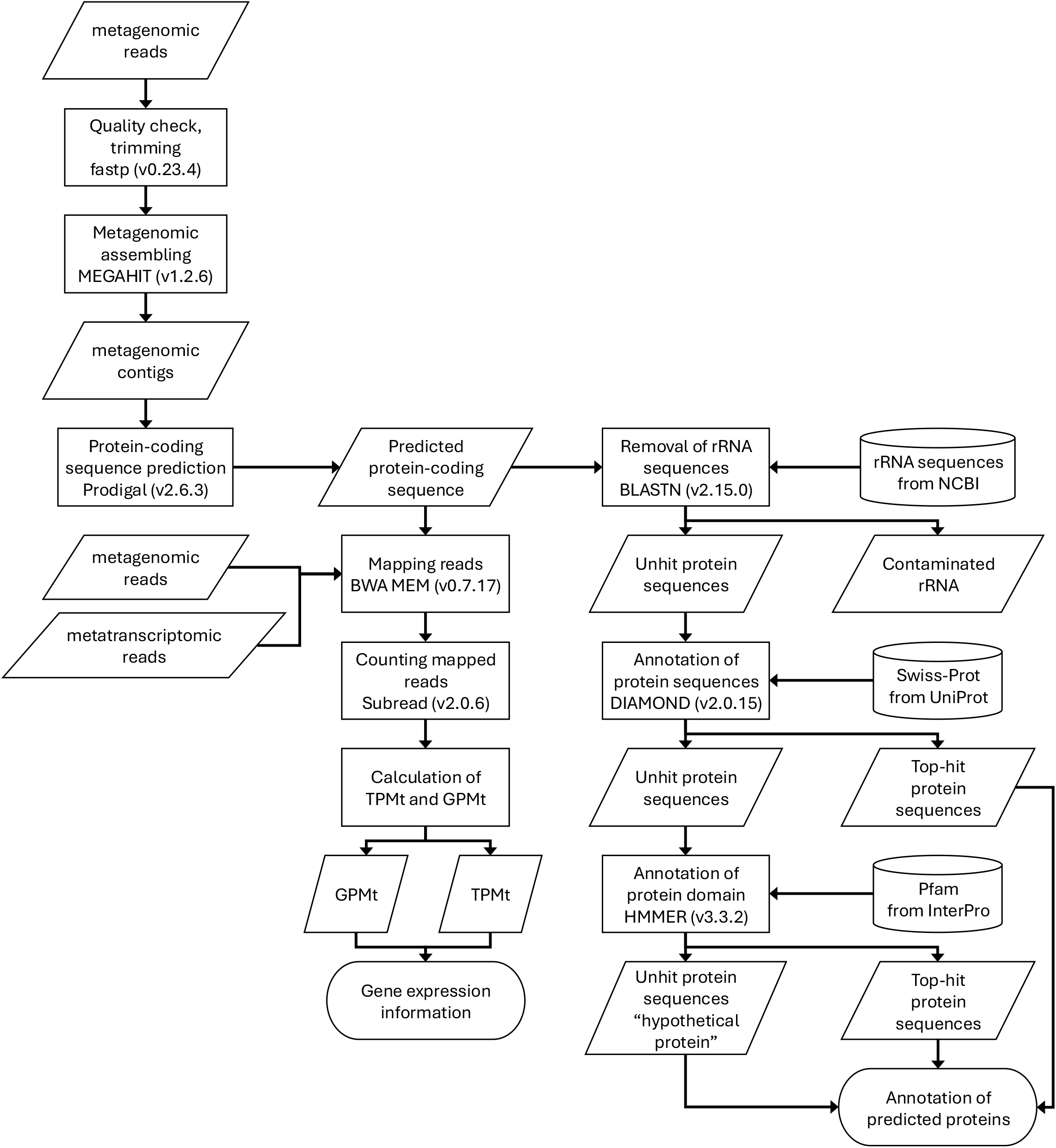
Data-driven analytical workflow for gene expression analysis in complex microbiomes.

## Results and Discussion

### Evaluation of contigs and mapping rates

The analytical workflow, encompassing processes from contig construction to gene expression quantification, was developed and integrated (Fig. 1). The following sections outline the considerations behind this workflow and present the results of gene expression analysis performed using the proposed framework.

To identify optimal reference sequences for gene expression analysis in complex microbiomes, three types of contigs were constructed from metagenomic and metatranscriptomic reads. The quality of these contigs was assessed, revealing that contigs assembled from metatranscriptomic reads using Trinity generally exhibited smaller N50 values compared to metagenomic contigs. While MAGs displayed improved N50 values relative to metagenomic contigs, their total length was often less than 10% of the metagenomic contigs. This trend was consistent across all analyzed samples (Supplementary Table 2).

Metagenomic and metatranscriptomic reads were then mapped to the three types of candidate reference sequences using two mapping tools, BWA MEM and Bowtie2. Across all samples, BWA MEM consistently achieved higher mapping rates than Bowtie2, regardless of the contig or read type (Fig. 2). This finding held true even when Bowtie2 was switched from end-to-end mode to local mode (data not shown). Notably, previous studies have reported superior mapping rates for BWA MEM in metagenomic samples [43].

**Figure 2.**
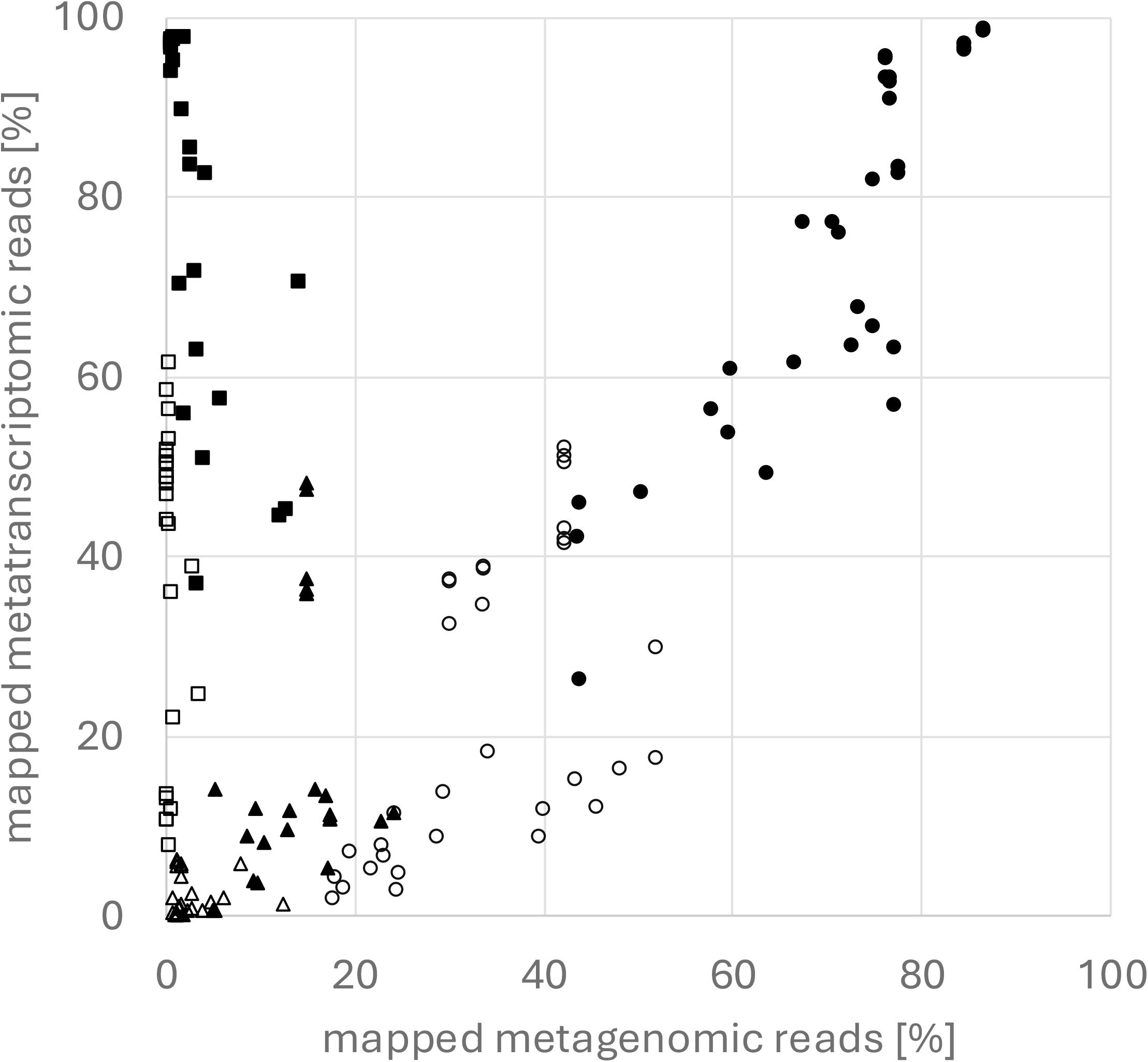
Mapping rates of metagenomic and metatranscriptomic reads. Reads were mapped using BWA-MEM (filled shapes) or Bowtie2 (outlined shapes) to protein-coding sequences derived from metagenomic contigs (circles), MAGs (triangles), or metatranscriptomic contigs (squares).

The mapping performance of the candidate reference sequences was further evaluated (Fig. 2). MAGs exhibited low mapping rates across all samples, reflecting their smaller sizes. Metatranscriptomic contigs demonstrated high mapping rates for metatranscriptomic reads but low rates for metagenomic reads. In contrast, metagenomic contigs yielded high mapping rates for both metagenomic and metatranscriptomic reads. This discrepancy is likely because metatranscriptomic contigs represent only expressed genes, whereas metagenomic contigs encompass both expressed and non-expressed genes. Therefore, metagenomic contigs are considered the optimal reference sequences, and BWA MEM the most suitable mapping tool, for comprehensive mapping of metagenomic and metatranscriptomic reads.

### Gene annotation

Predicted protein sequences derived from the metagenomic contigs were found to include rRNA sequences due to chance translational frames. To address this, BLASTN searches were conducted against rRNA sequence databases from NCBI. For instance, in the metagenome contigs assembled from SRR24888308, only 0.06% of the predicted protein-coding sequences were identified as rRNA. However, analysis of mapped read counts revealed that 0.15% (98,099 of 64,119,316) of metagenomic reads and 32.1% (23,574,758 of 73,419,607) of metatranscriptomic reads mapped to rRNA sequences. Since rRNA contamination can affect TPM calculations in gene expression analysis, these sequences were excluded before downstream analyses.

Although *in vitro* rRNA removal was performed in the referenced study [14], residual rRNA contamination persisted in metatranscriptomic reads, affecting predicted protein-coding sequences. Tools like SortMeRNA [44] can remove rRNA *in silico* from metatranscriptomic reads, potentially improving analytical accuracy. Future studies should address this issue further to ensure precise gene expression quantification.

Functional annotations were assigned to predicted protein sequences using the Swiss-Prot and Pfam databases. DIAMOND and HMMER were employed with an E-value threshold of 0.1 to achieve broad annotation coverage. Among 3,743,227 predicted protein-coding sequences derived from the metagenomic reads of SRR24888308, DIAMOND identified hits for 1,951,703 sequences (52.1%), HMMER for 838,155 sequences (22.4%), and 953,369 sequences (25.5%) were classified as hypothetical proteins.

The annotation results were consistent with ratios observed in fully sequenced bacterial genomes. For example, in *Bacillus subtilis* subsp. *subtilis* strain 168 (Genome assembly ASM904v112) [45], 13.3% of sequences are annotated as hypothetical proteins or pseudogenes, while in *Pseudomonas aeruginosa* PAO1 (Genome assembly ASM676v113) [46], this proportion rises to 40.7%. These results indicate that the annotation coverage achieved by the workflow is comparable to that of well-characterized genomes.

### Gene expression quantification

The preceding sections described the development of the workflow. This section presents the results of gene expression analysis conducted using the workflow.

In this analytical framework, gene expression levels were quantified using TPMt and GPMt. During contig construction from short-read data, variations in contig lengths were observed, even among contigs annotated as homologous genes. For example, acetyl-CoA hydrolase 1 (ACH1) coding sequences constructed from SRR24888635 ranged from 83 to 1,190 bp. Read counts mapped to contigs do not account for variations in predicted protein-coding sequence length, making them unsuitable for precise expression analysis. This limitation underscores the necessity of normalization methods, such as TPM and GPM, which adjust for sequence length and facilitate reliable gene expression analysis. In GPMt calculations, one count was added to the read counts to enable the computation of TPMt/GPMt ratios. Summing TPMt values for protein-coding sequences annotated as homologous genes provides an estimate of overall gene expression levels in the microbiome. However, when expression levels differ between samples, it is unclear whether gene expression per microbe has changed. To address this, TPMt/GPMt ratios were employed to assess gene expression intensity per microbe and identify genes that are actively expressed or not expressed. This approach provides a more nuanced evaluation of gene expression dynamics within microbial communities.

Gene expression analysis was performed using six metagenomic and six metatranscriptomic read datasets, and the results were compared with data from the reference study [14], focusing on specific target genes. In the reference study, gene expression levels were evaluated by summing read counts mapped to contigs annotated with the same Gene Ontology (GO) term. Similarly, in this study, TPMt values for predicted protein-coding sequences annotated as homologous genes were summed. The analysis revealed that *ACH1* was highly expressed in SRR24887267 and SRR24887404 samples, while acetone carboxylase beta subunit (*acxA*) and acetone carboxylase alpha subunit (*acxB*) were highly expressed in the SRR24887221 sample (Fig. 3D–F). These trends were consistent with the reference study [14] (Fig. 3A– C). Additionally, as reported in the reference study, no significant differences in metagenomic gene quantification values between samples were observed when GPMt values were summed without adding 1 to the read count (Fig. 3G–I). For example, while *acxA* exhibited a 171.57-fold difference in TPM values between SRR24887221 and SRR24887272, the GPM values differed by only 1.15-fold, suggesting that even samples with low TPM values retain the potential for gene expression.

**Figure 3.**
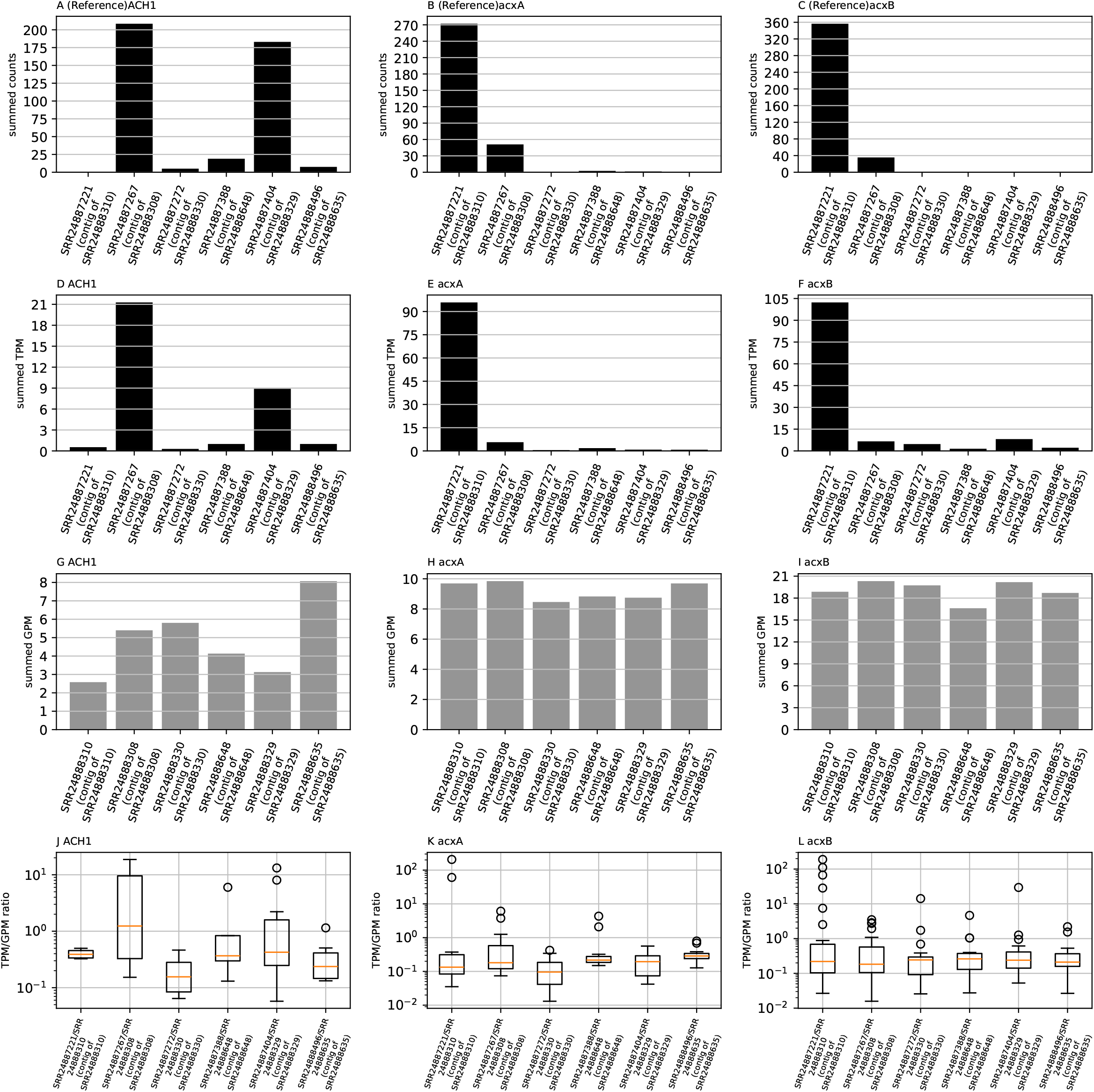
Gene expression analysis metrics. Panels A–C show summed read counts for each gene in the reference article, D–F display summed TPM values, G–I present summed GPM values, and J–L illustrate TPM/GPM ratios calculated in this study. Genes analyzed include *ACH1* (A, D, G, J), *acxA* (B, E, H, K), and *acxB* (C, F, I, L).

Using data not included in the reference study [14], expression intensities normalized by GPMt were visualized with boxplots. While *ACH1* displayed consistent expression patterns across analyses, *acxA* and *acxB* exhibited similar expression intensities across samples, with two outliers for *acxA* and six for *acxB* at higher expression levels (Fig. 3J–L). These findings suggest that *ACH1* may be regulated consistently across microorganisms, while microorganisms harboring *acxA* and *acxB* with outlier expression intensities may have distinct regulatory mechanisms. Variations in acetone carboxylase expression among species likely reflect these regulatory differences [47].

By analyzing metagenomic and metatranscriptomic reads using metagenomic contigs as reference sequences, this study successfully evaluated gene abundance and transcriptional activity in complex microbiomes. Additionally, it revealed the presence of genes with differential regulatory mechanisms among microorganisms. These findings support the conclusion that metagenomic contigs are more suitable than metatranscriptomic contigs or MAGs for gene expression analysis in complex microbiomes, as previously reported [10–12].

However, challenges remain in making comparisons among samples. Although gene expression analysis is based on reads mapped to predicted protein sequences with the same annotations, reference sequences differ across samples because metagenomic contigs are constructed individually for each sample. This limits the precise evaluation of gene expression variations between samples. Future work will focus on constructing unified reference sequences across samples to enable more accurate and consistent gene expression analysis in complex microbiomes.

Additionally, this workflow relies on relative quantification methods, such as GPM and TPM, to normalize abundance within each sample. Improving cross-sample quantification accuracy may require approaches such as incorporating internal standards during NGS read acquisition [48]. Furthermore, more efficient methods for extracting DNA and RNA from environmental microbiome samples need to be explored [49–51]. Combining such advanced sample preparation techniques with this workflow could further enhance the precision and scalability of quantitative gene expression analysis in complex microbiomes.

## Supplementary materials

**Supplementary Table 1**. List of 56 run IDs of 24 samples collected from Sequence Read Archive (SRA).

**Supplementary Table 2**. The quality of MAG, metagenomic and metatranscriptomic reads.

## Author contribution

Ryo Mameda designed the study, constructed and evaluated the workflow, and wrote the manuscript. Hidemasa Bono provided critical feedback and helped to shape the manuscript.

## Conflicts of interest

None declared.

## Funding

This research was supported by the Center of Innovation for Bio-Digital Transformation (BioDX), the open innovation platform for industry-academia co-creation (COI-NEXT), the Japan Science and Technology Agency (JST; Grant Number JPMJPF2010).

## Data availability

The workflow, comprising several scripts, is available in the GitHub repository at https://github.com/RyoMameda/workflow.

## Acknowledgment

Computations were partially conducted on the NIG supercomputer at the ROIS National Institute of Genetics.

## Notes

https://github.com/RyoMameda/workflow

### Competing Interest Statement

The authors have declared no competing interest.

## References

1. Sato Y, Hori T, Koike H et al. Transcriptome analysis of activated sludge microbiomes reveals an unexpected role of minority nitrifiers in carbon metabolism. Commun Biol 2019;2;:179 10.1038/s42003-019-0418-2.

2. Yang S, Zhang W, Yang B et al. Metagenomic evidence for antibiotic-associated actinomycetes in the Karamay Gobi region. Front Microbiol 2024;15;:1330880 10.3389/fmicb.2024.1330880.

3. Nelson MB, Martiny AC, Martiny JBH. Global biogeography of microbial nitrogen-cycling traits in soil. Proc Natl Acad Sci USA 2016;113;:8033–40 10.1073/pnas.1601070113.

4. Hartman WH, Ye R, Horwath WR et al. A genomic perspective on stoichiometric regulation of soil carbon cycling. The ISME Journal 2017;11;:2652–65 10.1038/ismej.2017.115.

5. Albertsen M, Hugenholtz P, Skarshewski A et al. Genome sequences of rare, uncultured bacteria obtained by differential coverage binning of multiple metagenomes. Nat Biotechnol 2013;31;:533–8 10.1038/nbt.2579.

6. Lasken RS, McLean JS. Recent advances in genomic DNA sequencing of microbial species from single cells. Nat Rev Genet 2014;15;:577–84 10.1038/nrg3785.

7. The University of Queenland. the Genome Taxonomy Database., https://gtdb.ecogenomic.org/ (29 December 2024, xdate last accessed).

8. Zhang Z, Wang J, Wang J et al. Estimate of the sequenced proportion of the global prokaryotic genome. Microbiome 2020;8;:134 10.1186/s40168-020-00903-z.

9. Yu K, Zhang T. Metagenomic and Metatranscriptomic Analysis of Microbial Community Structure and Gene Expression of Activated Sludge. Liles MR (ed.). PLoS ONE 2012;7;:e38183 10.1371/journal.pone.0038183.

10. Martinez X, Pozuelo M, Pascal V et al. MetaTrans: an open-source pipeline for metatranscriptomics. Sci Rep 2016;6;:26447 10.1038/srep26447.

11. Westreich ST, Treiber ML, Mills DA et al. SAMSA2: a standalone metatranscriptome analysis pipeline. BMC Bioinformatics 2018;19;:175 10.1186/s12859-018-2189-z.

12. Saenz C, Nigro E, Gunalan V et al. MIntO: A Modular and Scalable Pipeline For Microbiome Metagenomic and Metatranscriptomic Data Integration. Front Bioinform 2022;2;:846922 10.3389/fbinf.2022.846922.

13. Meziti A, Rodriguez-R LM, Hatt JK et al. The Reliability of Metagenome-Assembled Genomes (MAGs) in Representing Natural Populations: Insights from Comparing MAGs against Isolate Genomes Derived from the Same Fecal Sample. McBain AJ (ed.). Appl Environ Microbiol 2021;87;:e02593–20 10.1128/AEM.02593-20.

14. Honeker LK, Pugliese G, Ingrisch J et al. Drought re-routes soil microbial carbon metabolism towards emission of volatile metabolites in an artificial tropical rainforest. Nat Microbiol 2023;8;:1480–94 10.1038/s41564-023-01432-9.

15. Mendes LW, Raaijmakers JM, De Hollander M et al. Impact of the fungal pathogen Fusarium oxysporum on the taxonomic and functional diversity of the common bean root microbiome. Environmental Microbiome 2023;18;:68 10.1186/s40793-023-00524-7.

16. Freeman EC, Emilson EJS, Dittmar T et al. Universal microbial reworking of dissolved organic matter along environmental gradients. Nat Commun 2024;15;:187 10.1038/s41467-023-44431-4.

17. Li X, Bei Q, Rabiei Nematabad M et al. Time-shifted expression of acetoclastic and methylotrophic methanogenesis by a single Methanosarcina genomospecies predominates the methanogen dynamics in Philippine rice field soil. Microbiome 2024;12;:39 10.1186/s40168-023-01739-z.

18. SRA Toolkit Development Team. SRA Toolkit., https://github.com/ncbi/sra-tools (29 December 2024, xdate last accessed).

19. Chen S. Ultrafast one-pass FASTQ data preprocessing, quality control, and deduplication using fastp. iMeta 2023;2;:e107 10.1002/imt2.107.

20. Li D, Liu C-M, Luo R et al. MEGAHIT: an ultra-fast single-node solution for large and complex metagenomics assembly via succinct de Bruijn graph. Bioinformatics 2015;31;:1674–6 10.1093/bioinformatics/btv033.

21. Li D, Luo R, Liu C-M et al. MEGAHIT v1.0: A fast and scalable metagenome assembler driven by advanced methodologies and community practices. Methods 2016;102;:3–11 10.1016/j.ymeth.2016.02.020.

22. Kang DD, Li F, Kirton E et al. MetaBAT 2: an adaptive binning algorithm for robust and efficient genome reconstruction from metagenome assemblies. PeerJ 2019;7;:e7359 10.7717/peerj.7359.

23. Wu Y-W, Simmons BA, Singer SW. MaxBin 2.0: an automated binning algorithm to recover genomes from multiple metagenomic datasets. Bioinformatics 2016;32;:605–7 10.1093/bioinformatics/btv638.

24. Parks DH, Imelfort M, Skennerton CT et al. CheckM: assessing the quality of microbial genomes recovered from isolates, single cells, and metagenomes. Genome Res 2015;25;:1043–55 10.1101/gr.186072.114.

25. Matsen FA, Kodner RB, Armbrust EV. pplacer: linear time maximum-likelihood and Bayesian phylogenetic placement of sequences onto a fixed reference tree. BMC Bioinformatics 2010;11;:538 10.1186/1471-2105-11-538.

26. Hyatt D, Chen G-L, LoCascio PF et al. Prodigal: prokaryotic gene recognition and translation initiation site identification. BMC Bioinformatics 2010;11;:119 10.1186/1471-2105-11-119.

27. Grabherr MG, Haas BJ, Yassour M et al. Full-length transcriptome assembly from RNA-Seq data without a reference genome. Nat Biotechnol 2011;29;:644–52 10.1038/nbt.1883.

28. Shen W, Le S, Li Y et al. SeqKit: A Cross-Platform and Ultrafast Toolkit for FASTA/Q File Manipulation. Zou Q (ed.). PLoS ONE 2016;11;:e0163962 10.1371/journal.pone.0163962.

29. Shen W, Sipos B, Zhao L. SeqKit2: A Swiss army knife for sequence and alignment processing. iMeta 2024;3;:e191 10.1002/imt2.191.

30. Li H. Aligning sequence reads, clone sequences and assembly contigs with BWA-MEM. 2013 10.48550/arXiv.1303.3997.

31. Langmead B, Trapnell C, Pop M et al. Ultrafast and memory-efficient alignment of short DNA sequences to the human genome. Genome Biol 2009;10;:R25 10.1186/gb-2009-10-3-r25.

32. Langmead B, Salzberg SL. Fast gapped-read alignment with Bowtie 2. Nat Methods 2012;9;:357–9 10.1038/nmeth.1923.

33. Langmead B, Wilks C, Antonescu V et al. Scaling read aligners to hundreds of threads on general-purpose processors. Hancock J (ed.). Bioinformatics 2019;35;:421–32 10.1093/bioinformatics/bty648.

34. Danecek P, Bonfield JK, Liddle J et al. Twelve years of SAMtools and BCFtools. GigaScience 2021;10;:giab008 10.1093/gigascience/giab008.

35. NCBI. rRNA sequences., https://ftp.ncbi.nlm.nih.gov/blast/db/16S_ribosomal_RNA.tar.gz,https://ftp.ncbi.nlm.nih.gov/blast/db/18S_fungal_sequences.tar.gz,https://ftp.ncbi.nlm.nih.gov/blast/db/28S_fungal_sequences.tar.gz,https://ftp.ncbi.nlm.nih.gov/blast/db/LSU_prokaryote_rRNA.tar.gz,https://ftp.ncbi.nlm.nih.gov/blast/db/SSU_eukaryote_rRNA.tar.gz,https://ftp.ncbi.nlm.nih.gov/blast/db/LSU_eukaryote_rRNA.tar.gz (29 December 2024, date last accessed).

36. Camacho C, Coulouris G, Avagyan V et al. BLAST+: architecture and applications. BMC Bioinformatics 2009;10;:421 10.1186/1471-2105-10-421.

37. UniProt - EMBL-EBI. Swiss-Prot database., https://ftp.uniprot.org/pub/databases/uniprot/current_release/knowledgebase/complete/uniprot_sprot.fasta.g (29 December 2024, date last accessed).

38. Buchfink B, Reuter K, Drost H-G. Sensitive protein alignments at tree-of-life scale using DIAMOND. Nat Methods 2021;18;:366–8 10.1038/s41592-021-01101-x.

39. InterPro - EMBL-EBI. Pfam database., https://ftp.ebi.ac.uk/pub/databases/Pfam/current_release/Pfam-A.hmm.gz (29 December 2024, date last accessed).

40. HMMER., http://hmmer.org/ (29 December 2024, xdate last accessed).

41. Tange, Ole. GNU Parallel 20230922 (‘Derna’). 2023, 10.5281/zenodo.8374296.

42. Liao Y, Smyth GK, Shi W. featureCounts: an efficient general purpose program for assigning sequence reads to genomic features. Bioinformatics 2014;30;:923–30 10.1093/bioinformatics/btt656.

43. Jaillard M, Tournoud M, Meynier F et al. Optimization of alignment-based methods for taxonomic binning of metagenomics reads. Bioinformatics 2016;32;:1779–87 10.1093/bioinformatics/btw040.

44. Kopylova E, Noé L, Touzet H. SortMeRNA: fast and accurate filtering of ribosomal RNAs in metatranscriptomic data. Bioinformatics 2012;28;:3211–7 10.1093/bioinformatics/bts611.

45. Barbe V, Cruveiller S, Kunst F et al. From a consortium sequence to a unified sequence: the Bacillus subtilis 168 reference genome a decade later. Microbiology 2009;155;:1758–75 10.1099/mic.0.027839-0.

46. Stover CK, Pham XQ, Erwin AL et al. Complete genome sequence of Pseudomonas aeruginosa PAO1, an opportunistic pathogen. Nature 2000;406;:959–64 10.1038/35023079.

47. Sluis MK, Larsen RA, Krum JG et al. Biochemical, Molecular, and Genetic Analyses of the Acetone Carboxylases from Xanthobacter autotrophicus Strain Py2 and Rhodobacter capsulatus Strain B10. J Bacteriol 2002;184;:2969–77 10.1128/JB.184.11.2969-2977.2002.

48. Hardwick SA, Chen WY, Wong T et al. Synthetic microbe communities provide internal reference standards for metagenome sequencing and analysis. Nat Commun 2018;9;:3096 10.1038/s41467-018-05555-0.

49. Mori H, Kato T, Ozawa H et al. Assessment of metagenomic workflows using a newly constructed human gut microbiome mock community. DNA Research 2023;30;:dsad010 10.1093/dnares/dsad010.

50. Thorn CE, Bergesch C, Joyce A et al. A robust, cost-effective method for DNA, RNA and protein co-extraction from soil, other complex microbiomes and pure cultures. Molecular Ecology Resources 2019;19;:439–55 10.1111/1755-0998.12979.

51. Shaffer JP, Marotz C, Belda-Ferre P et al. A Comparison of DNA/RNA Extraction Protocols for High-Throughput Sequencing of Microbial Communities. BioTechniques 2021;70;:149–59 10.2144/btn-2020-0153.

